# G_αq_ sensitizes TRPM8 to inhibition by PI(4,5)P_2_ depletion upon receptor activation

**DOI:** 10.1101/410878

**Authors:** Luyu Liu, Yevgen Yudin, Chifei Kang, Natalia Shirokova, Tibor Rohacs

## Abstract

Activation of G-protein coupled receptors (GPCRs) was proposed to inhibit the cold and menthol sensitive Transient Receptor Potential Melastatin 8 (TRPM8) channels via direct binding of G_αq_ to the channel. It is well documented that TRPM8 requires the plasma membrane phospholipid phosphatidylinositol 4,5-bisphosphate [PI(4,5)P_2_ or PIP_2_] for activity. It was claimed however that a decrease in cellular levels of this lipid does not contribute to channel inhibition upon receptor activation. Here we show that supplementing the whole cell patch pipette with PI(4,5)P_2_ reduced inhibition of TRPM8 by activation of G_αq_-coupled receptors in mouse dorsal root ganglion (DRG) neurons. Activation of the same receptors induced Phospholipase C (PLC) activation and decreased plasma membrane PI(4,5)P_2_ levels in these neurons. PI(4,5)P_2_ also reduced inhibition of TRPM8 by activation of heterologously expressed G_αq_-coupled muscarinic M1 receptors. Co-expression of a constitutively active G_αq_ protein that does not couple to PLC inhibited TRPM8 activity, and in cells expressing this protein decreasing PI(4,5)P_2_ levels using a voltage sensitive 5’-phosphatase induced a stronger inhibition of TRPM8 activity than in control cells. Our data indicate that PI(4,5)P_2_ depletion plays an important role in TRPM8 inhibition upon GPCR activation, and G_αq_ inhibits the channel by reducing its apparent affinity for PI(4,5)P_2_ and thus sensitizes the channel to inhibition by decreasing PI(4,5)P_2_ levels.

## INTRODUCTION

TRPM8 is a non-selective Ca^2+^ permeable cation channel activated by cold temperatures, as well as chemical agonists such as menthol, icilin, and WS12 (Almaraz et al., 2014). It is expressed in the primary sensory neurons of dorsal root ganglia (DRG) and trigeminal ganglia (TG), and mice lacking these channels display deficiencies in sensing moderately cold (Bautista et al., 2007; Colburn et al., 2007; Dhaka et al., 2007) as well as noxious cold temperatures (Knowlton et al., 2010). TRPM8 may also play a role in cold allodynia induced by nerve injury (Xing et al., 2007).

DRG neurons express receptors for various inflammatory mediators, such as bradykinin, ATP and prostaglandins. Most of those receptors couple to heterotrimeric G-proteins in the G_q_ family, and lead to activation of phospholipase C (PLC) (Rohacs, 2016). Activation of these G_q_-coupled receptors induces thermal hyperalgesia (Caterina et al., 2000), via mechanisms including sensitizing the heat- and capsaicin-activated Transient Receptor Potential Vanilloid (TRPV1) channels (Cesare and McNaughton, 1996; Tominaga et al., 2001) and inhibiting the cold-activated TRPM8 channels (Premkumar et al., 2005; Zhang et al., 2012).

The mechanism of the inhibition of TRPM8 by G_q_-coupled receptors is not fully understood. Protein Kinase C (PKC) has been proposed to be involved in sensitization of TRPV1 (Zhang et al., 2008), but its involvement in TRPM8 inhibition is controversial (Premkumar et al., 2005; Zhang et al., 2012). It was proposed that binding of G_αq_ to TRPM8 mediates inhibition of the channel (Zhang et al., 2012). Similar to most TRP channels (Rohacs, 2014), TRPM8 requires the membrane phospholipid phosphatidylinositol 4,5-bisphosphate [PI(4,5)P_2_ or PIP_2_] for activity (Liu and Qin, 2005; Rohacs et al., 2005; Daniels et al., 2009; Zakharian et al., 2010). Ca^2+^ influx through the channel was shown to activate a Ca^2+^ sensitive PLC isoform, and the resulting decrease in PI(4,5)P_2_ levels limits channel activity, leading to desensitization (Rohacs et al., 2005; Daniels et al., 2009; Yudin et al., 2011; Yudin et al., 2016). Stimulation of G_q_-coupled receptors leads to the activation of PLCβ isoforms, which hydrolyze PI(4,5)P_2_, but it was questioned whether this leads to a decrease in PI(4,5)P_2_ levels in DRG neurons (Liu et al., 2010), and it was argued that a decrease in PI(4,5)P_2_ levels does not play a role in TRPM8 channel inhibition (Zhang et al., 2012).

Here we revisited this question, and demonstrate that the decrease in PI(4,5)P_2_ levels plays an important role in inhibition of TRPM8 by G_q_-coupled receptor activation in DRG neurons. First we show that application of a cocktail of inflammatory mediators that activate G_q_-coupled receptors decreased PI(4,5)P_2_ levels and inhibited TRPM8 activity in DRG neurons. Supplementing the whole cell patch pipette with PI(4,5)P_2_ alleviated the inhibition of TRPM8 by these receptor agonists. Excess PI(4,5)P_2_ also decreased the inhibition of heterologously expressed TRPM8 by activation of muscarinic M1 receptors. Co-expression of a constitutively active G_αq_ that is deficient in PLC activation inhibited TRPM8 activity, which is consistent with earlier results (Zhang et al., 2012). Co-expression of G_αq_ also increased the sensitivity of TRPM8 inhibition by decreasing PI(4,5)P_2_ levels using a specific voltage sensitive phosphoinositide 5-phosphatase, providing a mechanistic framework of how G_αq_ inhibits TRPM8. Overall, our data indicate that a decrease in PI(4,5)P_2_ levels and direct binding of G_αq_ converge on inhibiting TRPM8 activity upon cell surface receptor activation.

## MATERIALS AND METHODS

### DRG neuron isolation and preparation

All animal procedures were approved by the Institutional Animal Care and Use Committee at Rutgers New Jersey Medical School. Dorsal root ganglion (DRG) neurons were isolated from 2- to 6-month-old wild-type C57BL6 or TRPM8-GFP mice (Takashima et al., 2007) as described previously (Yudin et al., 2016) with some modifications. DRG neurons were isolated from mice of either sex anesthetized and perfused via the left ventricle with ice-cold Hank’s buffered salt solution (HBSS; Invitrogen). DRGs were harvested from all spinal segments after laminectomy and removal of the spinal column and maintained in ice-cold HBSS for the duration of the isolation. Isolated ganglia were cleaned from excess dorsal root nerve tissue and incubated in an HBSS-based enzyme solution containing 3 mg/ml type I collagenase (Worthington) and 5 mg/ml Dispase (Sigma) at 37°C for 35 min, followed by mechanical trituration by repetitive pipetting through an uncut 1000 µl pipette tip. Digestive enzymes were then removed after centrifugation of the cells at 80 × g for 10 min. Cells were then either directly re-suspended in growth medium and seeded onto glass coverslips coated with a mixture of poly-D-lysine (Invitrogen) and laminin (Sigma), or were first transfected using the Amaxa nucleoporator according to manufacturer’s instructions (Lonza Walkersville). Briefly, 60,000–100,000 cells obtained from each animal were re-suspended in 100 µl nucleofector solution or certain amount of transduction mix after complete removal of digestive enzymes. The cDNA used for transfecting neurons was prepared using the Endo-Free Plasmid Maxi Kit from QIAGEN. Vectors and reagent for transduction with the Green PIP_2_ Sensor (CAG-Promoter) #D0405G were obtained from Montana Molecular. Before seeding onto glass coverslips, cells were transduced according to the manufacturers instructions. Neurons were maintained in culture for at least 24 h before measurements in DMEM-F12 supplemented with 10% FBS (Thermo Scientific), 100 IU/ml penicillin and 100 µg/ml streptomycin.

### HEK cell culture and preparation

Human embryonic kidney 293 (HEK293) cells were obtained from the American Type Culture Collection (ATCC) (catalogue number CRL-1573) and were cultured in minimal essential medium (Invitrogen) containing supplements of 10% (v/v) Hyclone-characterized FBS (Thermo Scientific), 100 IU/ml penicillin, and 100 µg/ml streptomycin. Transient transfection was performed at ∼70% cell confluence with the Effectene reagent (QIAGEN) according to the manufacturer’s protocol. Cells were incubated with the lipid–DNA complexes overnight (16-20 h). The cDNA used for transfecting HEK cells was prepared using the Endo-Free Plasmid Maxi Kit from QIAGEN. 3G_qiq_ cDNA was obtained from Xuming Zhang’s laboratory (Zhang et al., 2012). 3G_qiq_Q209L was generated using the QuickChange Mutagenesis Kit (Agilent). Transduction was performed at ∼70% cell confluence with BacMam green PIP_2_ sensor with CAG promoter and BacMam M1 receptor (Montana Molecular) according to manufacturer’s protocol. Cells were incubated with transduction mix overnight (16-24 h). We then trypsinized and re-plated the cells onto poly-D-lysine-coated glass coverslips and incubated them for an additional at least 2 h (in the absence of the transfection reagent) before measurement. All mammalian cells were kept in a humidity-controlled tissue-culture incubator maintaining 5% CO_2_ at 37°C.

### Electrophysiology

Whole-cell patch-clamp recordings were performed at room temperature (27– 30°C) as described previously (Yudin et al., 2016). Patch pipettes were pulled from borosilicate glass capillaries (1.75 mm outer diameter, Sutter Instruments) on a P-97 pipette puller (Sutter Instrument) to a resistance of 4–6 MΩ. After formation of seals, the whole-cell configuration was established and currents were measured at a holding potential of −60 mV or with a voltage ramp protocol from -100 mV to +100 mV using an Axopatch 200B amplifier (Molecular Devices). Currents were filtered at 2 kHz using the low-pass Bessel filter of the amplifier and digitized using a Digidata 1440 unit (Molecular Devices). In some experiments, membrane potential pulses of indicated lengths to 100 mV were applied to activate the voltage-sensitive phosphatase (Ci-VSP), as described earlier (Velisetty et al., 2016). Measurements were conducted in solutions based on either Ca^2+^-free (NCF) medium containing the following (in mM): 137 NaCl, 4 KCl, 1 MgCl_2_, 5 HEPES, 5 MES, 10 glucose, 5 EGTA, pH adjusted to 7.4 with NaOH, or 2 mM Ca^2+^ (NCF) medium containing the following (in mM): 137 NaCl, 5 KCl, 1 MgCl_2_, 10 HEPES, 10 glucose, 2 CaCl_2_, pH adjusted to 7.4 with NaOH (Lukacs et al., 2013a). Intracellular solutions for DRG measurements (NIC-DRG) consisted of the following (in mM): 130 K-Gluconate, 10 KCl, 2 MgCl_2_, 2 Na_2_ATP, 0.2 Na_2_GTP, 1.5 CaCl_2_, 2.5 EGTA, 10 HEPES, pH adjusted to 7.25 with KOH. Intracellular solutions for HEK293 whole-cell measurements (NIC-HEK) consisted of the following (in mM): 130 KCl, 10 KOH, 3 MgCl_2_, 2 Na_2_ATP, 0.2 Na_2_GTP, 0.2 CaCl_2_, 2.5 EGTA, 10 HEPES, pH adjusted to 7.25 with KOH (Lukacs et al., 2013b). DiC_8_ PI(4,5)P_2_ was dissolved in NIC.

### Ca^2+^ imaging

Ca^2+^ imaging measurements were performed with an Olympus IX-51 inverted microscope equipped with a DeltaRAM excitation light source (Photon Technology International). DRG neurons or HEK cells were loaded with 1 µM fura-2 AM (Invitrogen) for 50 min before the measurement at room temperature (27–30°C), and dual-excitation images at 340 nm and 380 nm were recorded with a Roper Cool-Snap digital CCD camera. Measurements were conducted in NCF solution supplemented with 2 mM CaCl_2_. Image analysis was performed using the Image Master software (PTI).

### Confocal fluorescence imaging

Confocal measurements were conducted with an Olympus FluoView-1000 confocal microscope operation in frame scan mode (Olympus) at room temperature (∼25°C). GFP and YFP fluorescence (both excitation at 473 nm, emission at 520 nm) was recorded. DRG neurons or HEK cells were transfected with YFP-Tubby R332H-pcDNA3.1 or YFP-Tubby WT-pcDNA3.1 at least 24 hours before confocal measurements. Before each experiment, neurons or HEK cells were serum-deprived in Ca^2+^ free or 2 mM Ca^2+^-containing NCF solution for at least 20 min. For transduction experiments, DRG neurons or HEK cells were transduced at least 24 hours before confocal measurements by BacMam green PI(4,5)P_2_ sensor with CAG promoter and BacMam M1 receptor. Before each experiment, neurons or HEK cells were serum-deprived in DPBS (Sigma-Aldrich) for 30 min in the dark. Image analysis was performed using Olympus FluoView-1000 and ImageJ.

### Experimental Design and Statistical analysis

Data analysis was performed in Excel and Microcal Origin. Data collection was randomized. Data were analyzed with t-test, or Analysis of variance, p values are reported in the figures. Data are plotted as mean +/-standard error of the mean (SEM) for most experiments.

## RESULTS

### PI(4,5)P_2_ alleviates inhibition of TRPM8 activity by G_q_-coupled receptors in DRG neurons

Mouse DRG neurons express a number of different G_αq_-protein coupled receptors (Thakur et al., 2014). To identify receptors co-expressed with TRPM8, first we performed Ca^2+^ imaging experiments in DRG and TG neurons isolated from TRPM8-GFP reporter mice. We consecutively applied 500 µM menthol, 2 µM bradykinin, 1 µM prostaglandin E2 (PGE2), and 500 nM endothelin (ET) (Fig. S1 A, B). Most DRG neurons (62%) and TG neurons (84%) activated by menthol responded to either bradykinin, PGE2 or ET, but no individual agonist stimulated all TRPM8 positive neurons.

Next we measured Ca^2+^ signals in DRG neurons in response to the more specific TRPM8 agonist WS12 and to activate various G_αq_-coupled receptors we applied an inflammatory cocktail containing 100 µM ADP, 100 µM UTP, 500 nM bradykinin, 100 µM histamine, 100 µM serotonin, 10 µM PGE2 and 10 µM prostaglandin I2 (PGI2) (Fig. S1 C, D), similar to that used by (Zhang et al., 2012). We found that 11% of all DRG neurons were GFP positive and 57% of GFP-positive neurons responded to WS12 (Fig. S1 E). Within these GFP positive neurons, almost all neurons activated by WS12 responded to the inflammatory cocktail. Only one neuron (out of 214 neurons) was stimulated by WS12 but not by the inflammatory cocktail (Fig. S1 C, D, E).

Next we preformed whole-cell patch clamp experiments on TRPM8-GFP reporter mouse DRG neurons and tested the effect of supplementing the patch pipette solution with the water-soluble diC_8_ PI(4,5)P_2_. We used a protocol of consecutive 1 min applications of 10 µM WS12 in Ca^2+^-free NCF solution. Inflammatory cocktail was applied 1 min before the third application of WS12 (Fig. 1 A, B). Without diC_8_ PI(4,5)P_2_, current amplitudes induced by WS12 with inflammatory cocktail were reduced to 47% of currents induced by WS12 only (Fig. 1 A,C,D). When we included diC_8_ PI(4,5)P_2_ in the patch pipette solution, current amplitudes induced by WS12 with inflammatory cocktail were reduced only to 91% of currents induced by WS12 only (Fig. 1 B,C,D). These findings show TRPM8 channel activity is inhibited by G_αq_-protein coupled receptors in DRG neurons, and this inhibition is alleviated by excess PI(4,5)P_2_.

**Figure 1.**
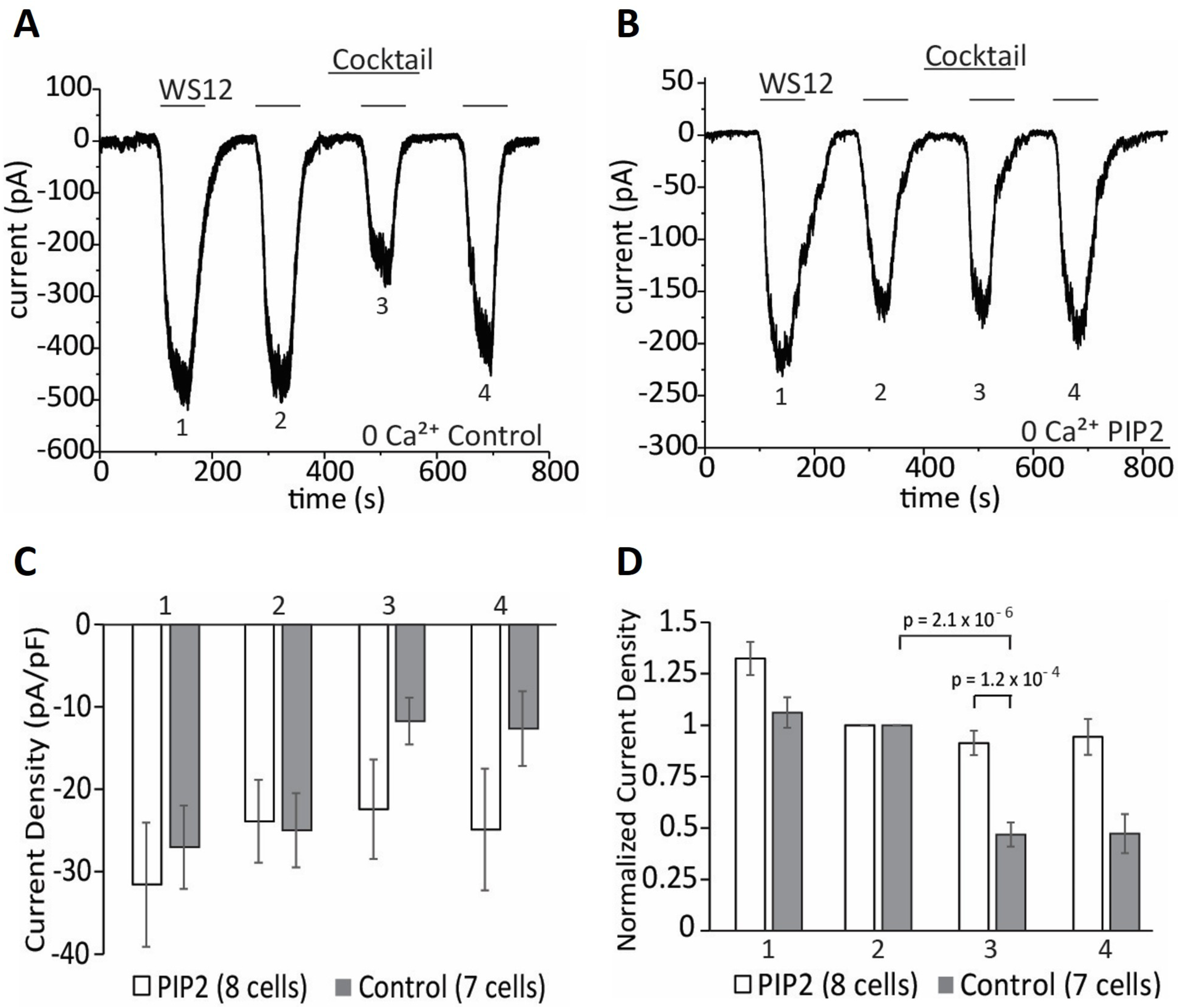
Intracellular dialysis of PI(4,5)P_2_ alleviates inhibition of TRPM8 activity by G_q_-coupled receptors in DRG neurons. ***A, B*,** Representative whole-cell voltage-clamp traces of inward currents recorded at -60 mV from isolated DRG neurons. Measurements were conducted in Ca^2+^-free NCF solution. Control-no lipid supplement (***A***), 100 µM diC_8_ PI(4,5)P_2_ supplement (***B***). 10 µM WS12 was applied to activate TRPM8 channels for 4 times. Cocktail containing 100 µM ADP, 100 µM UTP, 500 nM bradykinin, 100 µM histamine, 100 µM serotonin, 10 µM PGE2, 10 µM PGI2 was applied to activate various G_q_-coupled receptors. ***C, D*,** Statistical analysis of n=7 neurons (control group) and n=8 neurons (diC_8_ PI(4,5)P_2_ group) displayed. Bars represent mean ± SEM; statistical significance was calculated with two-way analysis of variance. Four peaks of current responses in each group were analyzed. Current values were normalized to the peak of the second current response in each group in ***D***.

### Receptor-induced decrease of PI(4,5)P_2_ in the plasma membrane

There are conflicting result on whether plasma membrane PI(4,5)P_2_ levels decrease in DRG neurons in response to stimulating endogenous G_q_-coupled bradykinin receptors (Liu et al., 2010; Lukacs et al., 2013b), and there is no information on the effects of other G_q_-coupled receptors (Rohacs, 2016). Therefore we tested if there is a decrease in PI(4,5)P_2_ levels in response to the inflammatory cocktail we used in our electrophysiology experiments. We transfected DRG neurons with the YFP-tagged R322H Tubby PI(4,5)P_2_ sensor using the Amaxa nucleoporator system. This YFP-tagged phosphoinositide binding domain of the Tubby protein contains the R322H mutation that decreases its affinity for PI(4,5)P_2_, thus increasing its sensitivity to small, physiological decreases in PI(4,5)P_2_ levels (Quinn et al., 2008; Lukacs et al., 2013b). The Tubby-based fluorescent sensor binds to PI(4,5)P_2_ specifically; it is located in the plasma membrane in resting conditions; when PI(4,5)P_2_ levels decrease, this indicator is translocated into the cytoplasm, which we monitored using confocal microscopy. Application of the inflammatory cocktail induced rapid translocation of the fluorescent sensor from the plasma membrane to the cytoplasm in the neuron cell body (Fig. 2 A, B, middle panels) as well as in the neuronal processes (Fig. 2 A, B, bottom panels), indicating decreased PI(4,5)P_2_ levels. Figs. 2 C-E summarize these results.

**Figure 2.**
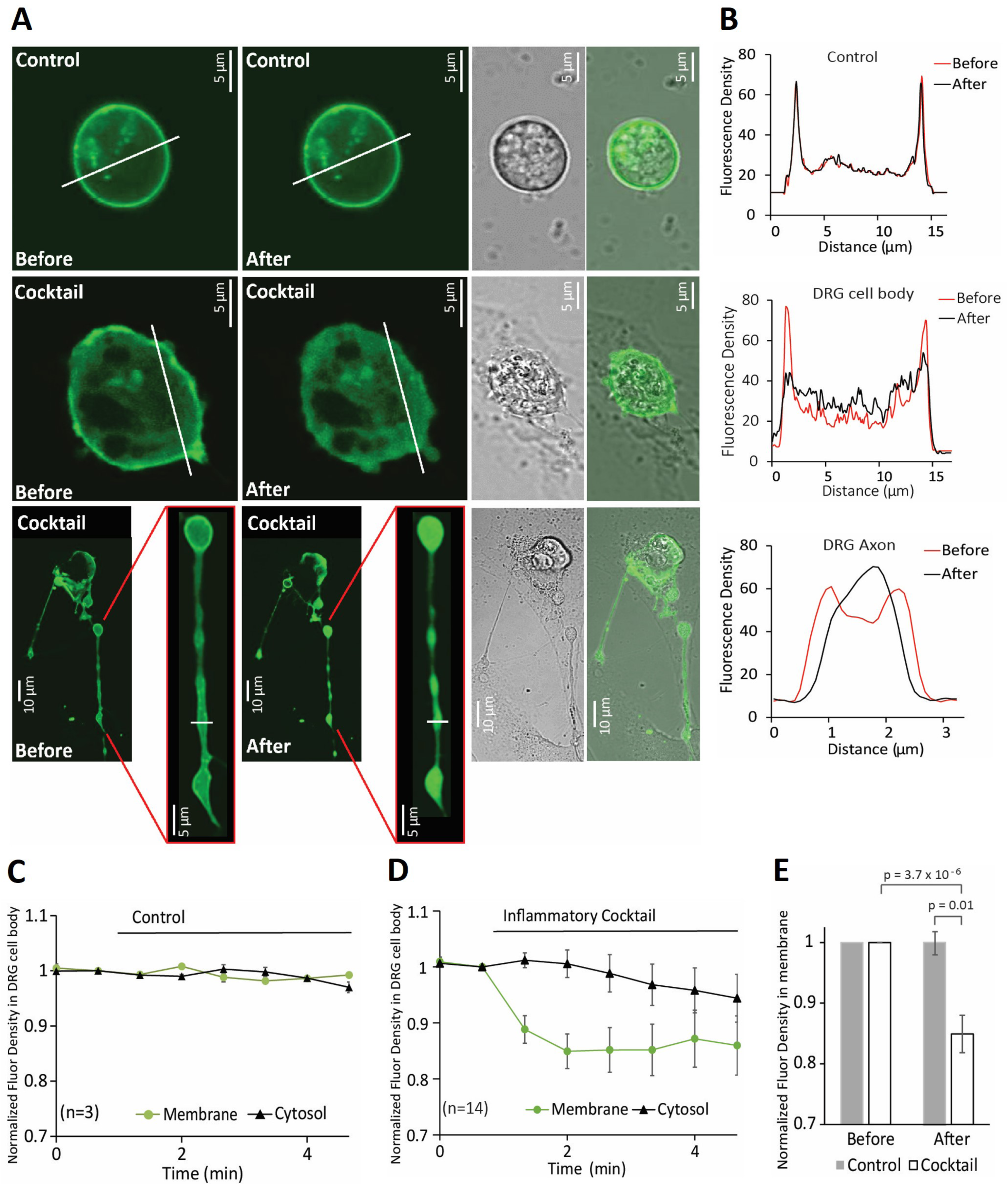
Receptor-induced decrease of PI(4,5)P_2_ in the plasma membrane of DRG neurons. ***A,*** Confocal images of wild type mouse DRG neurons transfected with YFP-tagged R322H Tubby domain as reporters of plasma membrane PI(4,5)P_2_. Images are representatives of reporter distribution before and after application of either inflammatory cocktail or control solution (2 mM Ca^2+^ NCF). Measurements were conducted in 2 mM Ca^2+^ NCF solution. Neurons were serum-deprived in 2 mM Ca^2+^ NCF solution for 20 min. ***B,*** Graphs correspond to fluorescence intensities plotted along the lines indicated in each image in ***A*** before (red) and after (black) application of cocktail or control. ***C, D,*** Time courses of fluorescence density changes for YFP-tagged R322H Tubby in response to control solution (***C***) or inflammatory cocktail (***D***) in plasma membrane and cytosol of neuron cell body. ***E,*** Statistical analysis of fluorescence density right before and 1 min after application of inflammatory cocktail (n=14) and control solution (n=3) displayed. Bars represent mean ± SEM; statistical significance was calculated with two-way analysis of variance. Fluorescence density values were normalized to the time point of right before application of inflammatory cocktail or control solution for each group.

We also monitored PI(4,5)P_2_ levels using a different fluorescence based sensor the BacMam PI(4,5)P_2_ sensor kit from Montana Molecular. This green PI(4,5)P_2_ sensor is based on a dimerization-dependent fluorescent PI(4,5)P_2_ binding protein (Tewson et al., 2013). The kit also contains cDNA for the M1 muscarinic receptor, a G_αq_-protein coupled receptor, to serve as a positive control (Thakur et al., 2014). Isolated neurons transduced with green BacMam PI(4,5)P_2_ sensor showed prominent plasma membrane labeling. Administration of inflammatory cocktail induced robust decrease of fluorescence levels in the plasma membrane, indicating decreased PI(4,5)P_2_ levels. Carbachol was applied at the end to activate M1 receptors, which induced a further decrease of fluorescence in most DRG neurons (Fig. 3 A-F).

**Figure 3.**
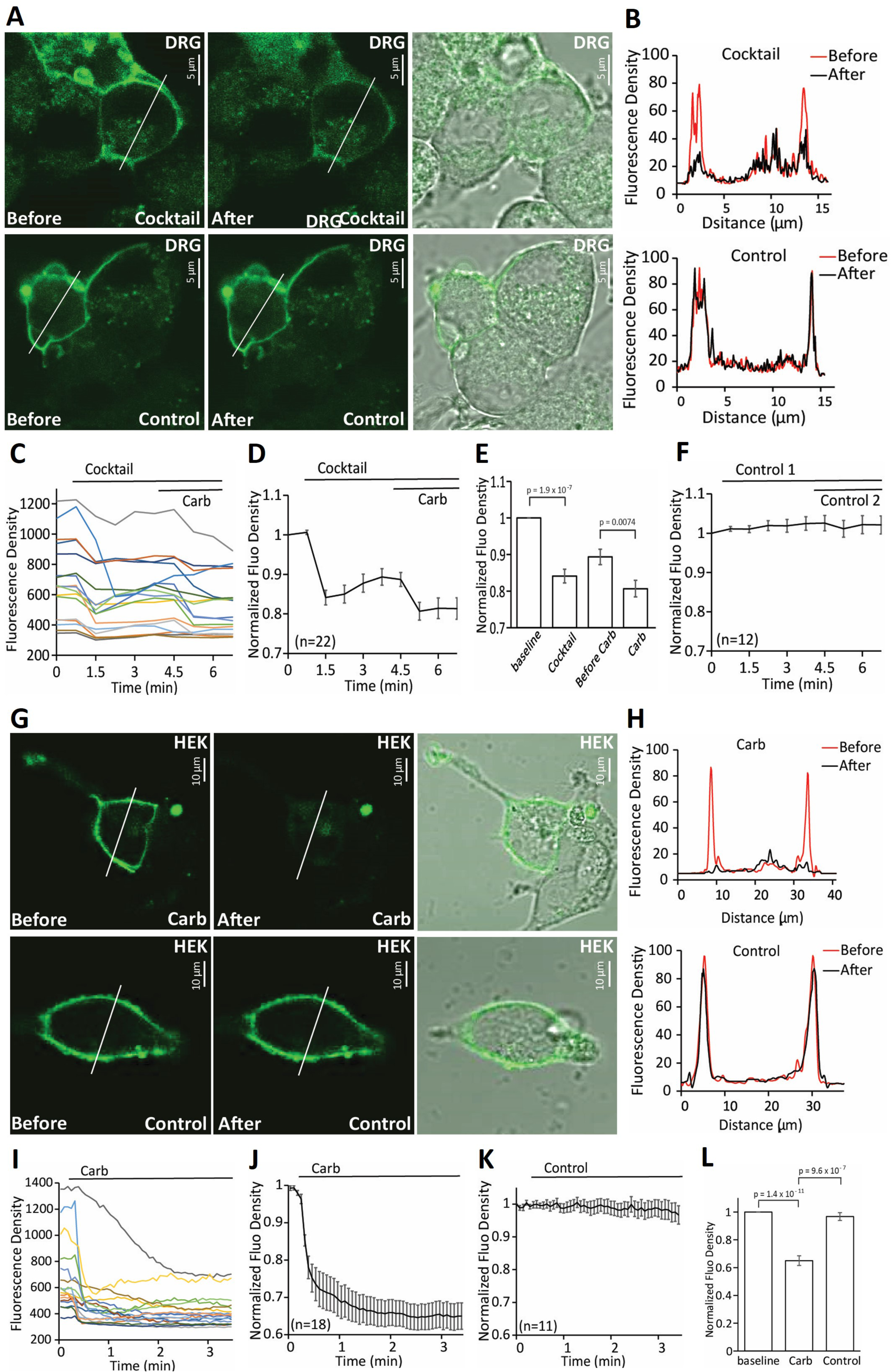
Receptor-induced decrease of PI(4,5)P_2_ in DRG neurons and HEK cells. ***A,*** Confocal images of wild type mouse DRG neurons tranducted with Green PI(4,5)P_2_ sensor BacMam and human muscarinic cholinergic receptor 1 (M1). Images are representatives of reporter distribution before and after application of either cocktail or control solution (2 mM Ca^2+^ NCF). Measurements were conducted in 2 mM Ca^2+^ NCF solution. ***B,*** Graphs correspond to fluorescence intensities plotted along the lines indicated in each image in ***A*** before (red) and after (black) application of cocktail or control. ***C,*** Representative traces of time course changes for Green PI(4,5)P_2_ sensor BacMam in response to cocktail followed by 100 µM carbachol in DRG neurons. ***D,*** Averaged time course of fluorescence density changes for Green PI(4,5)P_2_ sensor BacMam in response to cocktail followed by 100 µM carbachol in DRG neurons. **E,** Statistical analysis of fluorescence density at baseline, 40 seconds after application of cocktail, right before application of carbachol and 40 seconds after application of carbachol in neurons (n=22). Bars represent mean ± SEM; statistical significance was calculated with two-way analysis of variance. Fluorescence density values were normalized to baseline. ***F,*** Time course of fluorescence density changes for Green PI(4,5)P_2_ sensor BacMam in response to two applications of control solution (2 mM Ca^2+^ NCF) in DRG neurons (n=12). ***G,*** Confocal images of HEK cells transduced with Green PI(4,5)P_2_ sensor BacMam as reporters of PI(4,5)P_2_ and human muscarinic receptor 1 (M1) receptor. Images are representatives of reporter distribution before and after application of either 100 µM carbachol or control solution (2 mM Ca^2+^ NCF). Measurements were conducted in 2 mM Ca^2+^ NCF solution. ***H,*** Graphs correspond to fluorescence intensities plotted along the lines indicated in each image in ***G*** before (red) and after (black) application of 100 µM carbachol or control. ***I,*** Representative traces of time course changes for Green PI(4,5)P_2_ sensor BacMam in response to carbachol. ***J, K,*** Average time course of fluorescence density changes for Green PI(4,5)P_2_ sensor BacMam in response to carbachol or control (2 mM Ca^2+^ NCF) in HEK cells. **L,** Statistical analysis of fluorescence density at baseline and after application of carbachol (n=18) or control (n=11). Bars represent mean ± SEM; statistical significance was calculated with two-way analysis of variance. Fluorescence density values were normalized to baseline

We also tested receptor-induced depletion of PI(4,5)P_2_ in HEK293 cells transduced with the green BacMam PI(4,5)P_2_ sensor and BacMam M1 receptor. After 24-hour culture, HEK cells showed prominent plasma membrane labeling. Administration of carbachol in 2 mM Ca^2+^ NCF solution induced robust decrease of fluorescence levels in the plasma membrane, no changes were detected in “bleaching” control experiments (Fig. 3 G-L). Based on these results, we conclude that activation of G_q_-coupled receptors decreases PI(4,5)P_2_ levels in the plasma membrane.

### PI(4,5)P_2_ alleviates inhibition of TRPM8 activity by heterologously expressed G_q_-coupled receptors

We showed that PI(4,5)P_2_ alleviated inhibition of TRPM8 activity by G_q_-coupled receptors in DRG neurons. Next we tested if this was reproducible in HEK293 cells co-expressing TRPM8 and M1 receptors, which were reported to inhibit TRPM8 (Li and Zhang, 2013). To this end, we performed whole-cell patch clamp experiments with, or without supplementing the patch pipette solution with the water-soluble diC_8_ PI(4,5)P_2_ (Fig. 4 A, B). First we used a protocol of consecutive 20-second applications of 200 µM menthol in Ca^2+^-free NCF solution. Carbachol was applied 80 seconds before the fourth application of menthol. Current amplitudes induced by menthol were partially reduced by carbachol to 58% (at +60 mV) and 60% (at -60 mV) of currents induced by menthol only. When the patch pipette contained diC_8_ PI(4,5)P_2_, there was no decrease in current amplitudes after carbachol application (Fig. 4 D). We also tested whether protein kinase C (PKC) played a role in this inhibition. Previous studies showed controversial effects of PKC in TRPM8 inhibition induced by G_q_-coupled receptors (Premkumar et al., 2005; Zhang et al., 2012). We co-applied the PKC inhibitor BIM IV (1 µM) with 100 µM carbachol for 80 seconds before the fourth application of menthol (Fig. 4 C). We found no significant effect of BIM IV (Fig. 4 D), similar to the findings of (Zhang et al., 2012).

**Figure 4.**
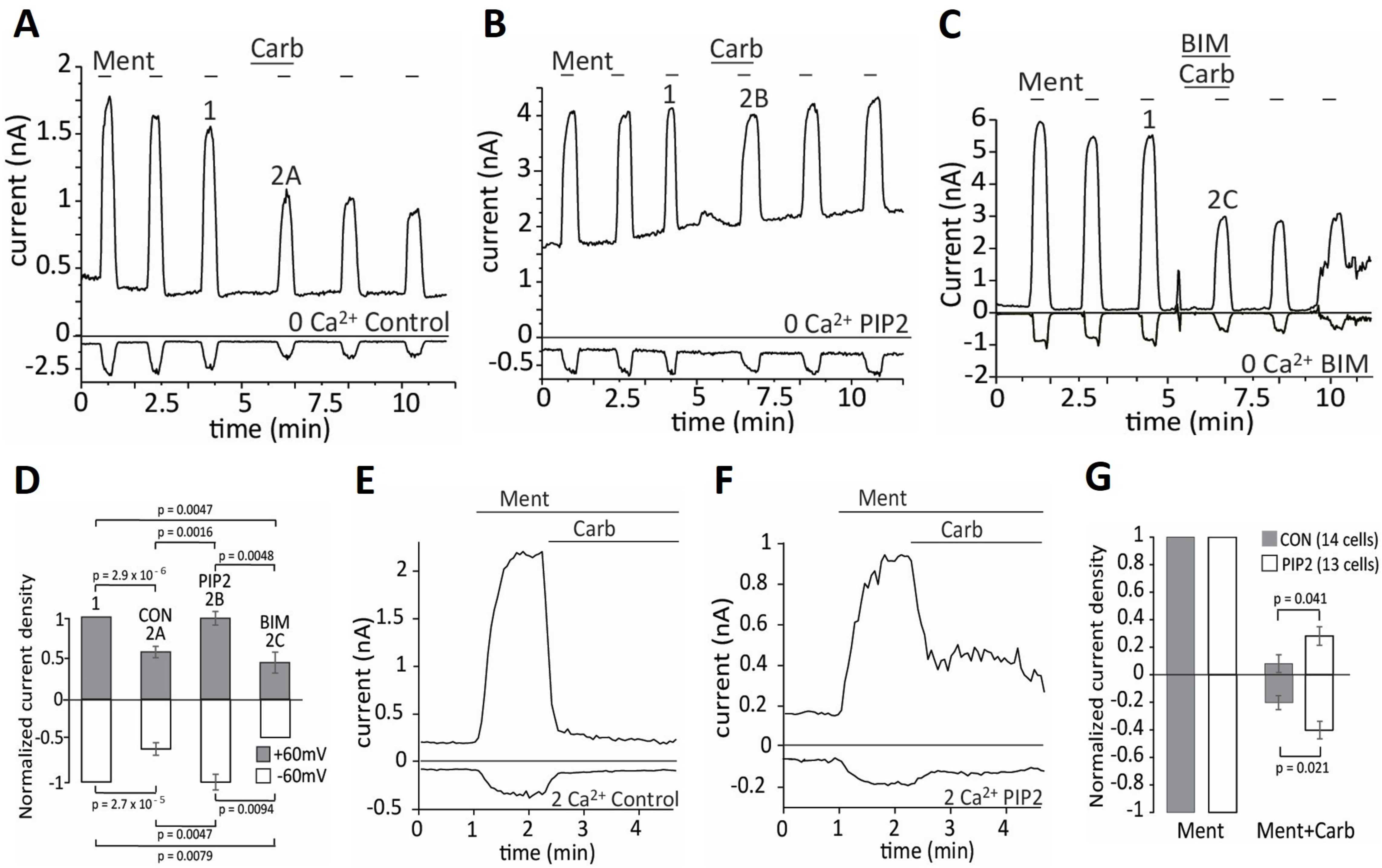
Intracellular dialysis of PI(4,5)P_2_ alleviates inhibition of TRPM8 activity by G_q_-coupled receptors in HEK cells. ***A, B*, C,** Representative whole-cell voltage-clamp traces of inward currents recorded at -60 mV and + 60 mV in Ca^2+^-free NCF solution for HEK cells transfected with TRPM8 channels and M1 receptors. Control-no lipid supplement (***A, C***), 100 µM diC_8_PI(4,5)P_2_ supplement (***B***). 200 µM Menthol was applied to activate TRPM8 channels for 6 times. 100 µM carbachol and 1 µM BIM were applied to activate M1 receptor or inhibit protein kinase C (***C***), respectively. ***D,*** Statistical analysis of n=12 neurons (control), n=7 (diC_8_ PI(4,5)P_2_) and n=4 neurons (BIM) both at -60 mV and + 60 mV (corresponding to panel ***A, B, C***) displayed. Bars represent mean ± SEM; statistical significance was calculated with two-way analysis of variance. Current values were normalized to the 3^rd^ current response to menthol (labeled as 1). The 4^th^ current response to menthol with co-application of carbachol was labeled as 2A, 2B, 2C in groups of control, diC_8_ PI(4,5)P_2_ and BIM separately. ***E, F,*** Representative whole-cell voltage-clamp traces of inward currents recorded at – 60 mV and + 60 mV in 2 mM Ca^2+^ NCF solution for HEK cells transfected with TRPM8 channels and M1 receptors; the applications of 200 µM Menthol and 100 µM carbachol were indicated by the horizontal lines; 100 µM diC_8_ PI(4,5)P_2_ was added into NIC in ***F***, no lipid solutions was added into NIC in ***E***. ***G,*** Statistical analysis of n=14 neurons (control) and n=13 (diC_8_ PI(4,5)P_2_) both at -60 mV and +60 mV (corresponding to panel ***E, F***) displayed. Bars represent mean ± SEM. *p<0.05 (two-sample Student’s *t* test). Current values at the time point of 1.5 min after application of carbachol were analyzed. Current values were normalized to the peak current value under menthol application before carbachol.

Next we performed similar experiments in NCF solution containing 2 mM Ca^2+^ (Fig. 4 E-G). Upon application of carbachol, TRPM8 currents induced by menthol were reduced to 18% at +60 mV and 29% at -60 mV. The inhibition was partially rescued by adding diC_8_ PI(4,5)P_2_ into patch pipette solution; current levels decreased to 37% (at +60 mV) and 46% (at -60 mV) under these conditions (Fig. 4 G).

Next we examined whether the enhanced inhibition in the presence of extracellular Ca^2+^ was because of different levels of PI(4,5)P_2_ depletion. We transfected HEK293 cells with the YFP-tagged wild-type Tubby and M1 receptor, and monitored translocation with confocal microscopy. The YFP-tagged wild type Tubby has higher affinity for PI(4,5)P_2_ than the YFP-tagged R322H Tubby (Quinn et al., 2008), therefore it may be more sensitive to changes at high levels of PI(4,5)P_2_ depletion. HEK293 cells transfected with YFP-tagged WT tubby showed prominent plasma membrane labeling and application of 100 µM carbachol both in 2 mM Ca^2+^ NCF solution and in Ca^2+^-free NCF solution induced translocation of fluorescent sensors from the plasma membrane to the cytoplasm (Fig. 5 A-E). Fluorescence levels in 2 mM Ca^2+^ NCF solution decreased more in plasma membrane and increased more in cytosol than in Ca^2+^-free NCF solution (Fig. 5C) and this difference was statistically significant (Fig. 5E). This indicates that M1 receptor activation induced a larger decrease in PI(4,5)P_2_ levels in the presence of extracellular Ca^2+^, compared to that in the absence of extracellular Ca^2+^. This correlates well with the stronger inhibition of TRPM8 in the presence of extracellular Ca^2+^. We also used lower affinity YFP-tagged R322H Tubby in 2 mM Ca^2+^ NCF solution and in Ca^2+^-free NCF solution and found that there was no significant difference of PI(4,5)P_2_ level changes detected (Fig. S2 A-E). This is likely due to the large decrease of PI(4,5)P_2_ levels in both conditions, differentiating between which is outside the dynamic range of this probe.

**Figure 5.**
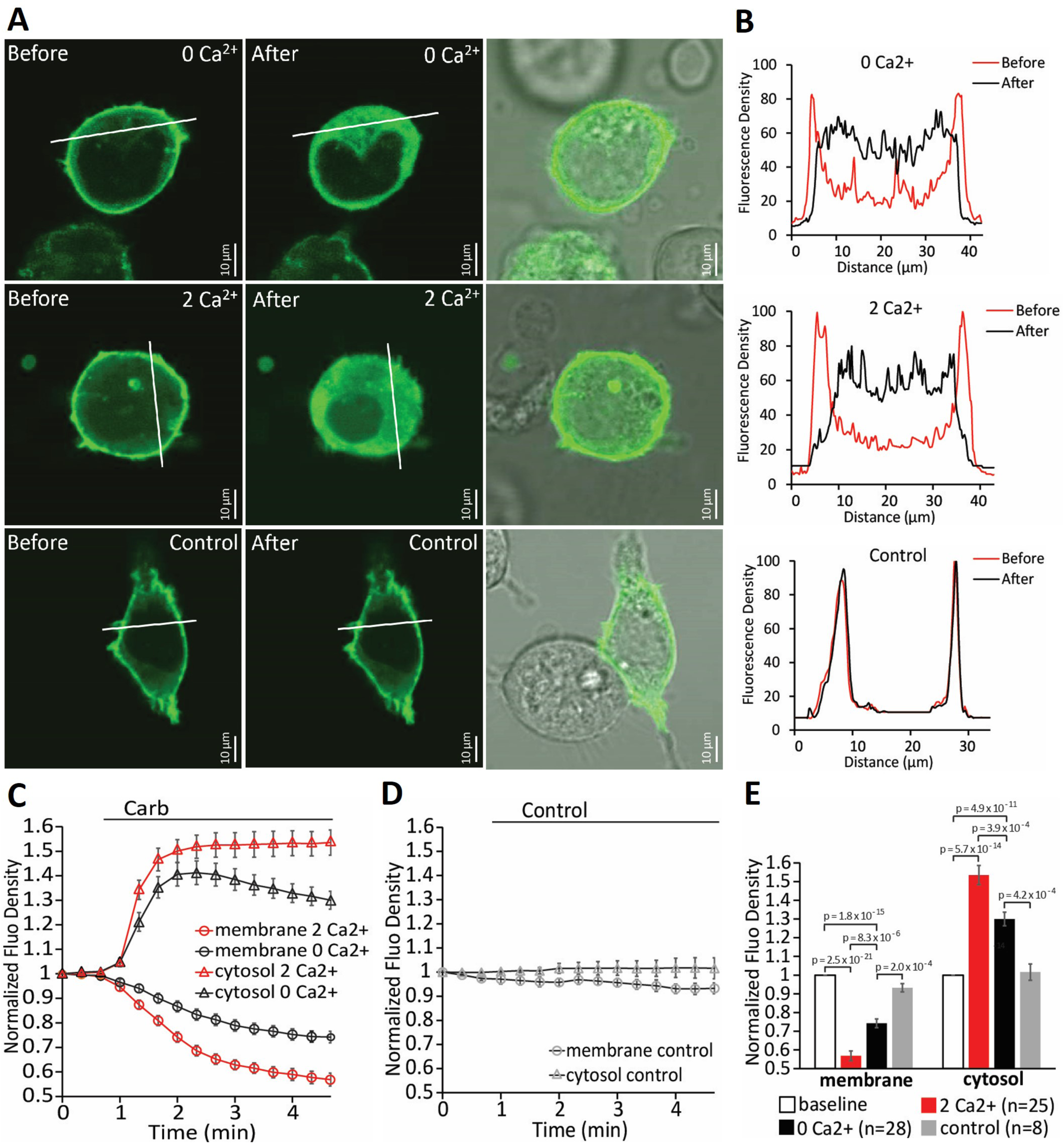
Receptor-induced depletion of PI(4,5)P_2_ in the plasma membrane of HEK cells in the presence or absence of calcium. ***A,*** Confocal images of HEK cells transfected with YFP-tagged wild type Tubby domain as reporters of plasma membrane phosphoinositides and M1 receptor in 2 mM Ca^2+^ NCF or Ca^2+^-free NCF solution. Images are representatives of reporter distribution before and after application of either 100 µM carbachol or control solution (2 mM Ca^2+^ NCF). ***B,*** Graphs correspond to fluorescence intensities plotted along the lines indicated in each image in ***A*** before (red) and after (black) application of carbachol or control. ***C, D,*** Time courses of fluorescence density changes for YFP-tagged WT Tubby in response to carbachol (***C***) or control (***D***) in plasma membrane and cytosol of HEK cells. ***E,*** Statistical analysis of fluorescence density at baseline and at the end after application of carbachol (n=25 for 2 mM Ca^2+^ NCF, n=28 for Ca^2+^-free NCF) and control (n=8) displayed. Bars represent mean ± SEM; statistical significance was calculated with two-way analysis of variance. Fluorescence density values were normalized to the baseline for each group.

### G_αq_ inhibits TRPM8 in the absence of PLC activation

G_αq_ is inactive in the GDP-bound form in resting cells, and active in the GTP-bound form upon receptor activation, when it binds to and activates phospholipase Cβ isoforms (PLCβ). 3G_qiq_ is a G_αq_ mutant that does not couple to PLC, and 3G_qiq_Q209L is a constitutive active Gαq mutant that also does not couple to PLC (Zhang et al., 2012). We used Fura2 AM to measure Ca^2+^ level changes upon menthol application in HEK293 cells co-expressed with TRPM8 and either 3G_qiq_ or 3G_qiq_Q209L (Fig. 6 A, B). Menthol-induced Ca^2+^ level increase was significantly less in cells with 3G_qiq_Q209L than in cells with wild 3G_qiq_ or TRPM8 only (Fig. 6 A-D). This is consistent with the whole-cell patch-clamp results of Zhang et al. 2012, who found that 3G_qiq_Q209L inhibits TRPM8 activity and 3G_qiq_ does not.

**Figure 6.**
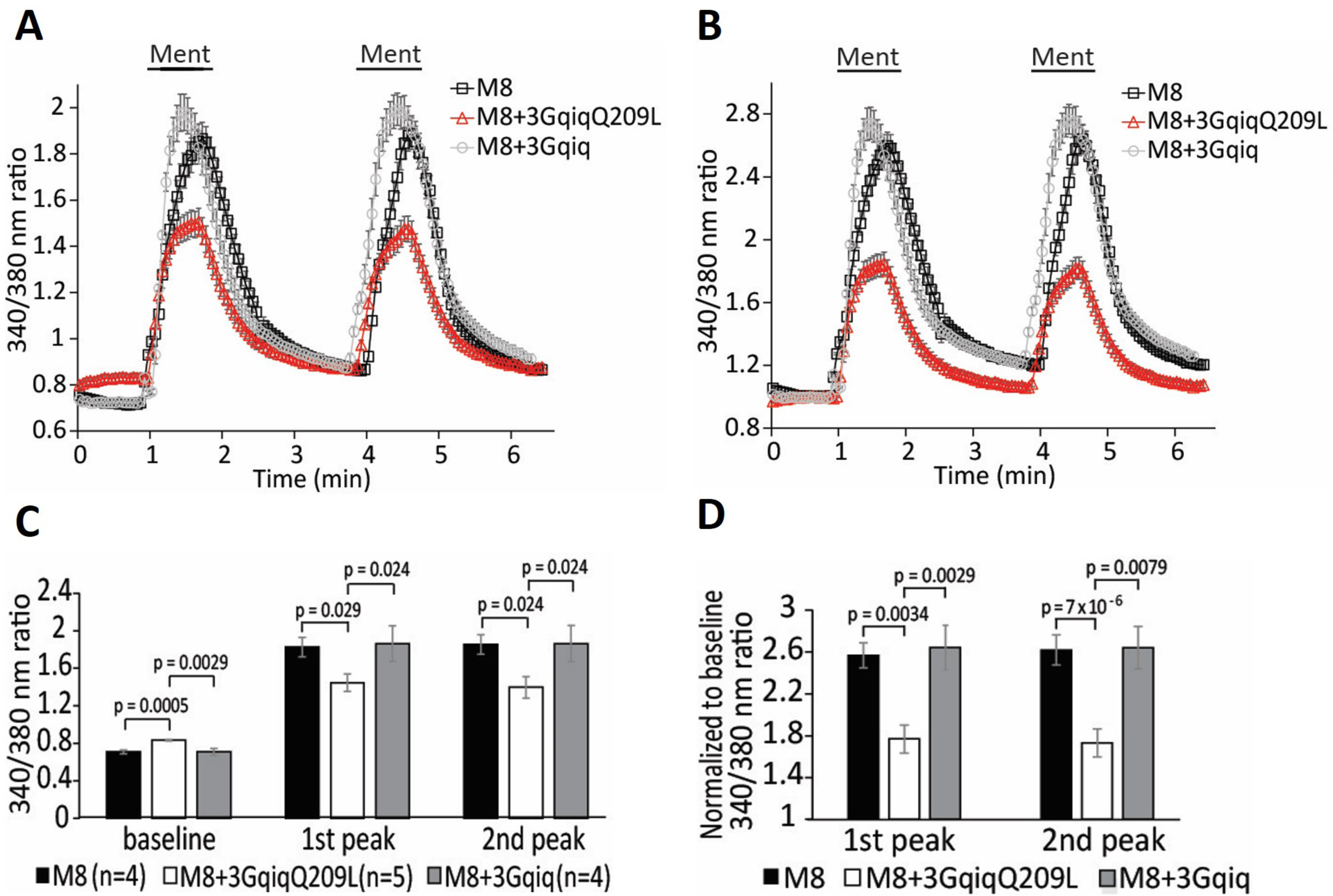
TRPM8 is inhibited by G_αq_ when PLC is not activated. ***A,*** Calcium imaging measurements (mean ± SEM) of HEK cells transfected with TRPM8, plus 3GqiqQ209L or 3Gqiq in response to two applications of 200 µM Menthol. ***B,*** Pooled data of ***A*** normalized to baseline. ***C, D,*** corresponds to ***A*** and ***B,*** separately. Bars represent mean ± SEM; statistical significance was calculated with two-way analysis of variance. Fluorescence values of 340/380 ratio were normalized to the baseline for each group. Two peaks of menthol responses were analyzed. TRPM8 group (n=4 slides, including 103 cells), TRPM8+3GqiqQ209L group (n=5 slides, including 84 cells), TRPM8 + 3Gqiq group (n=4 slides, including 103 cells)

### PI(4,5)P_2_ depletion inhibits TRPM8 more efficiently in the presence of G_αq_

To bypass the PLC pathway, we used a voltage-sensitive 5’ phosphatase (Ci-VSP) to deplete PI(4,5)P_2_ by converting it to PI(4)P (Iwasaki et al., 2008). Whole-cell patch-clamp experiments were performed in HEK293 cells co-transfected with TRPM8 and Ci-VSP, with or without 3G_qiq_Q209L in Ca^2+^-free NCF solution (Fig. 7 A-D). Ci-VSP was activated by depolarizing pulses to 100 mV for 0.1 s, 0.2 s, 0.3 s, and 1 s; menthol-induced currents were gradually reduced to 85%, 66%, 51%, and 32% respectively in cells without 3G_qiq_Q209L. In cells expressing 3G_qiq_Q209L, the same depolarizing pulses induced larger reductions in menthol-evoked currents, to 32%, 20%, 18%, and 15% respectively (Fig. 7 D). These results indicate that TRPM8 is inhibited more by PI(4,5)P_2_ depletion in the presence of G_αq_.

**Figure 7.**
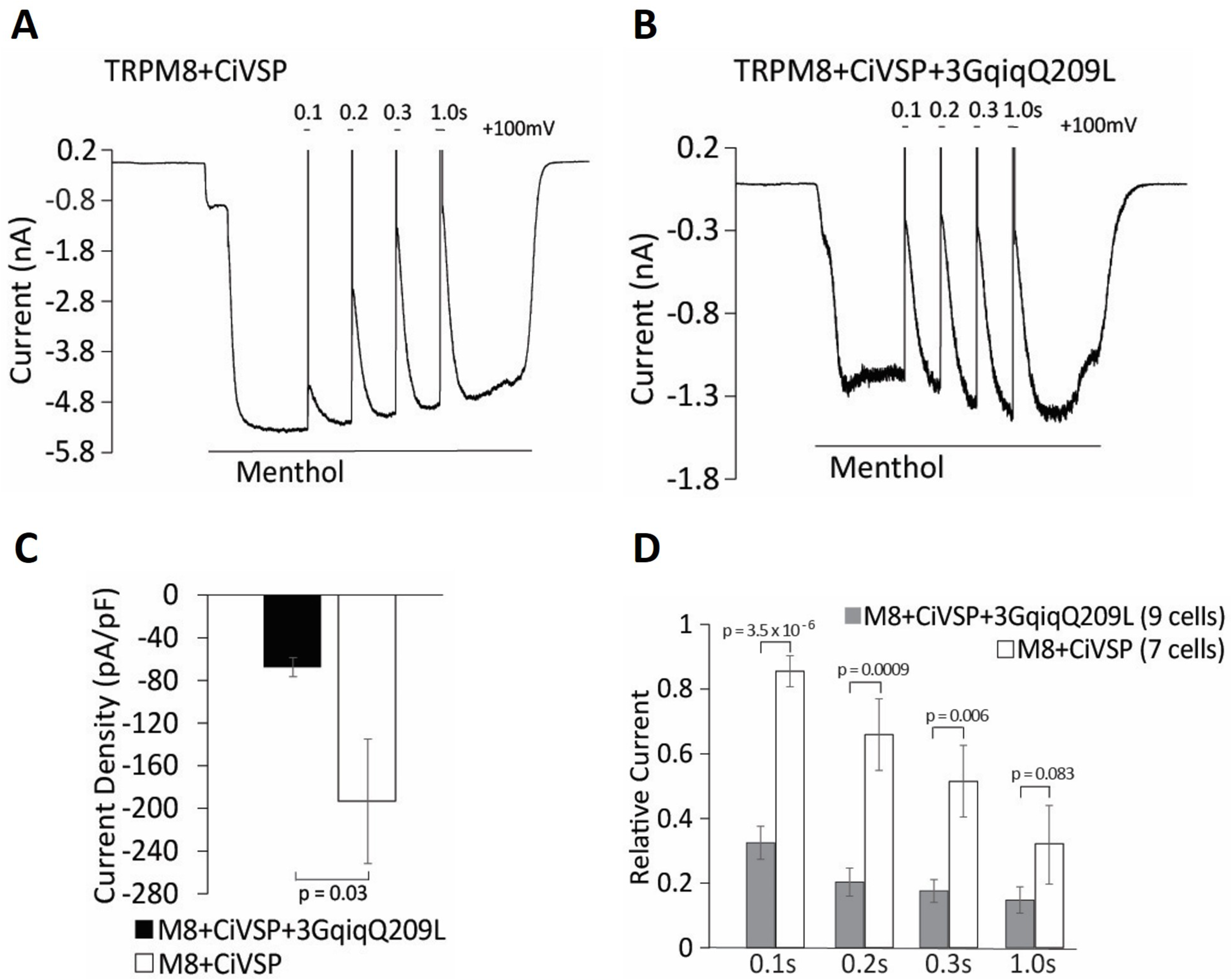
TRPM8 is inhibited more efficiently by PI(4,5)P_2_ depletion in the presence of G_αq_. ***A, B*,** Representative whole-cell voltage-clamp traces of inward currents recorded at -60 mV from HEK cells transfected with TRPM8 and the voltage sensitive phosphatase CiVSP (***A***), or with ciVSP, TRPM8 and 3GqiqQ209L (***B***). Measurements were conducted in Ca^2+^-free NCF solution. TRPM8 channels were activated by 500 µM menthol. CiVSP was activated by +100 mV steps for 0.1, 0.2, 0.3 and 1 second. ***C, D*,** Statistical analysis of ***A*** and ***B*** displayed. Bars represent mean ± SEM; statistical significance was calculated with two-sample Student’s *t* test, two-way analysis of variance. Current peak and current density upon voltage steps to +100 mV for 0.1, 0.2, 0.3 and 1 second were analyzed for each group (TRPM8 + CiVSP, n=7; TRPM8 + CiVSP + 3GqiqQ209L, n=9) in ***C***. Relative currents were analyzed upon voltage steps to +100 mV for 0.1, 0.2, 0.3 and 1 second in ***D***.

We conclude that PI(4,5)P_2_ alleviates inhibition of TRPM8 activity by G_q_-coupled receptors in DRG neurons. G_αq_ not only directly inhibits TRPM8 as proposed by Zhang et al 2012, but also sensitizes the channel to inhibition by decreasing PI(4,5)P_2_ levels, therefore the two pathways converge to inhibit TRPM8 activity upon GPCR activation.

## DISCUSSION

Converging evidence demonstrate that TRPM8 channels require the plasma membrane phospholipid PI(4,5)P_2_ for activity. TRPM8 currents show a characteristic decrease (rundown) in excised inside out patches, which can be restored by applying PI(4,5)P_2_ to the cytoplasmic surface of the membrane patch (Liu and Qin, 2005; Rohacs et al., 2005). Increasing endogenous PI(4,5)P_2_ levels with MgATP also restored TRPM8 activity in excised patches after current rundown (Yudin et al., 2011). PI(4,5)P_2_ is also required for the cold-or menthol-induced activity of the purified TRPM8 protein in planar lipid bilayers, indicating a direct effect on the channel (Zakharian et al., 2009; Zakharian et al., 2010). PI(4,5)P_2_ was also shown to modulate the cold threshold of the channel (Fujita et al., 2013). In accordance with dependence of the activity of TRPM8 on PI(4,5)P_2_, it was shown that selectively decreasing PI(4,5)P_2_ levels in intact cells is sufficient to inhibit channel activity either by using specific chemically inducible phosphoinositide phosphatases (Varnai et al., 2006; Daniels et al., 2009), or voltage sensitive phosphatases such as ciVSP (Yudin et al., 2011).

Stimulating cell surface receptors that couple to G_αq_ and activate PLC inhibits TRPM8 activity (Liu and Qin, 2005; Premkumar et al., 2005; Zhang et al., 2012; Li and Zhang, 2013; Than et al., 2013). Based on the following arguments it was proposed that this inhibition proceeds via direct binding of G_αq_ to the channel (Zhang et al., 2012). Application of G_αq_ and GTPγS inhibited TRPM8 activity in excised inside out patches in the presence of PI(4,5)P_2_. Direct binding of G_αq_ to TRPM8 was demonstrated by immunoprecipitation. Co-expression of a constitutively active mutant (Q209L) chimera between G_αi_ and G_αq_ (3G_αqiq_) that does not activate PLC inhibited TRPM8 activity (Zhang et al., 2012).

It was also proposed that decreasing PI(4,5)P_2_ levels do not contribute to TRPM8 inhibition by G_q_-coupled receptors (Zhang et al., 2012) based on the following data: Histamine inhibited TRPM8 in the presence of the PLC inhibitor U73122. The inhibition however was less that without the drug, indicating potential involvement of PLC activation. Two mutants in the putative TRP domain were also tested, and the authors argued that since these “PI(4,5)P_2_ insensitive” TRPM8 mutants were inhibited by GPCR activation, PI(4,5)P_2_ depletion is not involved. These mutants however are more sensitive to PI(4,5)P_2_ depletion, due to their decreased apparent affinity for the lipid (Rohacs et al., 2005), and indeed the K995Q mutant was somewhat more inhibited by histamine than wild-type TRPM8, and the extent of inhibition for the R1008Q is difficult to evaluate due to the very small current amplitudes (Zhang et al., 2012). Finally, the constitutively active Q209L mutant of the 3Gα_qiq_ chimera that does not activate PLC inhibited TRPM8 activity when co-expressed in HEK cells. The inhibition however was substantially smaller than that induced by the constitutively active G_αq_ indicating PLC dependent mechanisms may also involved (Zhang et al., 2012). In conclusion, most of the data discussed here is compatible with PI(4,5)P_2_ depletion contributing to TRPM8 inhibition.

To test the involvement of PI(4,5)P_2_ depletion, we supplemented the whole cell patch pipette with the lipid, and found that this maneuver alleviated inhibition of TRPM8 activity by G_q_-coupled receptor activation both in DRG neurons, and in an expression system. In DRG neurons, we used an inflammatory cocktail, similar to that used by (Zhang et al., 2012) for two reasons. First, DRG neurons are highly heterogenous, and we could not find a single agonist that reliably induced a Ca^2+^ signal indicating PLC activation in all TRPM8 positive neurons. Second, inflammation is mediated not by a single pro-inflammatory agonist, but rather a combination of them, which is sometime referred to as an “inflammatory soup”.

While activation of PLC induces PI(4,5)P_2_ hydrolysis, the extent to which PI(4,5)P_2_ is decreased is debated (Nasuhoglu et al., 2002), and is clearly cell type and agonist specific (van der Wal et al., 2001). Liu et al., using a fluorescent PI(4,5)P_2_ sensor, the YFP-tagged PI(4,5)P_2_ binding domain of the tubby protein (Quinn et al., 2008), showed that bradykinin activated PLC in rat DRG neurons, but it did not decrease in PI(4,5)P_2_ levels (Liu et al., 2010). Our earlier data using the same sensor show that bradykinin induced a small decrease in PI(4,5)P_2_ levels in mouse DRG neurons (Lukacs et al., 2013b). Here we demonstrate that application of the same inflammatory G_q_-activating cocktail we used in our patch clamp measurements decreased PI(4,5)P_2_ levels in a substantial portion of DRG neurons, using the tubby-based PI(4,5)P_2_ sensor. Note that this decrease was smaller than that induced by activating heterologously expressed M1 receptors therefore in native cells, additional mechanisms may be needed to induce substantial inhibition of TRPM8 activity.

We found that co-expressing the PLC deficient constitutively active 3Gα_qiq_-Q209L inhibited menthol-induced Ca^2+^ signals in HEK cells expressing TRPM8, confirming the results of (Zhang et al., 2012). We also found that the presence of 3Gα_qiq_-Q209L markedly increased the sensitivity of inhibition of TRPM8 by decreasing PI(4,5)P_2_ levels with the voltage sensitive phosphoinositide 5-phosphatase ciVSP. These results indicate the G_αq_ binding to the channel decreases the apparent affinity of the channel for PI(4,5)P_2_. It is quite likely that this decreased sensitivity to PI(4,5)P_2_ would render basal PI(4,5)P_2_ levels to be less efficient in maintaining channel activity, thus explaining the reduced TRPM8 current levels upon G_αq_ binding. This decreased apparent affinity would also allow even small decreases in PI(4,5)P_2_ concentrations, such as those occurring during physiological receptor activation, to inhibit channel activity.

Overall we conclude that upon receptor activation, direct binding of G_αq_ not only directly inhibits TRPM8, but also sensitizes it to PI(4,5)P_2_ depletion, and thus the two pathways converge and synergize in reducing channel activity.

## ACKNOWLEDGEMENTS

T.R. was supported by NIH grants NS055159 and GM093290. The authors thank Dr. Joshua Berlin for his insightful comments, Dr. David Julius (UCSF) for providing the TRPM8 clone, Dr. David McKemy (University of Southern California) for providing the GFP-TRPM8 mouse line, Dr. Yasushi Okamura (Osaka University, Japan) for providing the ci-VSP and dr-VSP clones, Drs. Nikita Gamper (University of Leeds) and Andrew Tinker (University College London) for providing the tubby-R332H-YFP clone, Dr. Xuming Zhang (Aston University, Birmingham, UK) for providing the 3G_qiq_ clone, and Linda Zabelka for maintaining the mouse colony.

